# Metabolomic comparison using *Streptomyces* spp. as a factory of secondary metabolites

**DOI:** 10.1101/2022.12.01.518800

**Authors:** Rene Flores Clavo, Alana Kelyene Pereira, Nataly Ruiz Quiñones, Jonas Henrique Costa, Taícia Pacheco Fill, Fabiana Fantinatti Garboggini

## Abstract

Understanding extremophiles and their usefulness in biotechnology involves studying their habitat, physiology and biochemical adaptations, as well as their ability to produce biocatalysts, in environments that are still poorly explored. In northwestern Peru, which has saline lagoons of marine origin Pacific Ocean, the other site is from the coast of Brazil of the Atlantic Ocean. Both environments are considered extreme. The objective of the present work was to compare two different strains isolated from these extreme environments at the metabolic level using molecular network methodology through the Global Natural Products Molecular Social Network (GNPS). In our study, the MS/MS spectra from the network were compared with GNPS spectral libraries, where the metabolites were annotated. Differences were observed in the molecular network presented in the two strains of *Streptomyces* spp. coming from these two different environments. Within the annotated compounds from marine bacteria, the metabolites characterized for *Streptomyces* sp. B-81 from Peruvian marshes were lobophorins A (1) and H (2), as well as divergolides A (3), B (4) and C (5). *Streptomyces* sp. 796.1 produced different compounds, such as glucopiericidin A (6) and dehy-dro-piericidin A1a (7). The search for new metabolites in underexplored environments may therefore reveal new metabolites with potential application in different areas of biotechnology.

## 1. Introduction

Approximately 22,500 biologically active substances are obtained from microorganisms, 45% of which are represented by Actinomycetes and 70% of which are *Streptomyces* metabolites; however, information on substances isolated from microorganisms inhabiting saline environments is scarce [1,2,3]. The search for these microorganisms has been mainly associated with the production of antibiotics and antitumor substances [4]. Halophilic and halotolerant strains show heterogeneous physiological characteristics for different genera because these bacteria can synthesize secondary metabolites to cope with the high salinity and extreme temperature conditions of their environments [5,6]. These extreme conditions favor the development of metabolic competitiveness for the production of enzymes, providing adaptation to the high salinity of the environment [4]. The production of bioactive molecules from actinobacteria other than *Streptomyces* allows extending the search for new molecules and interactions. During the last five years, constant isolation efforts of microorganisms from various environmental niches have resulted in the recovery of thousands of members of the actinobacteria affiliated with different taxonomic classes [5]. The members of the genus *Streptomyces* stand out as producers of the largest amount of secondary metabolites already described [6]. Various studies have revealed an approach combining LC–MS/MS and molecular networking as a rapid analytical method for the identification of compounds [7]; this microbial group has a unique ability to produce new products, mainly antibiotics [8,9].

[8] Investigated the metabolic profiling of a cultured endophyte strain (*Streptomyces* sp. HKI0576) by HPLC–MS and revealed a complex metabolome, *Streptomyces* sp. 12A35 was isolated from deep sea sediment collected from the South China Sea and showed promising antibacterial activities attributed to the spirorotronate antibiotics lobophorins B, F, H, and I from a marine origin [9]; Piericidins and Glucopiericidins A and B isolated from the culture broth of *Streptomyces pactum* S48727 [10], a new natural product of the lobophorin family designated lobophorin K, from cultures of the marine actinobacteria *Streptomyces* sp. M-207 [11]; A cryptic BGC for type I polyketides was activated by metabolic engineering methods, enabling the discovery of a known compound, lobophorin CR4, of deep-sea-derived *Streptomyces olivaceus* SCSIO T05 [12].

A study reported in Salar de Huasco in the Atacama Desert, considered a polyextreme environment in Chile, [13] described that species of the genus *Streptomyces* were dominant, and a preliminary study [14] present in the salt ponds of Bayovar and Morrope in Peru showed physicochemical characteristics common to extremophilic bacteria and ample potential to produce bioactive compounds. These extreme conditions encourage the development of metabolic competitiveness for the production, for example, of enzymes [15]. The application of various targeted studies on the biosynthesis of natural products allows a better understanding of the full potential of the producing microorganisms, thus increasing the return on investment in the search for new compounds [16].

In this study, we used a molecular networking approach for the rapid detection of molecules from two strains of *Streptomyces* spp. obtained from extreme environments: Saloons northwestern of Perú and Cabo Frio in Brazil (Figure 1). The detection method, based on UHPLC–MS/MS combined with data analysis using Global Natural Product Social Molecular Networking (GNPS), led to the annotation of very different compounds produced by each of the *Streptomyces* spp.

**Figure 1.**
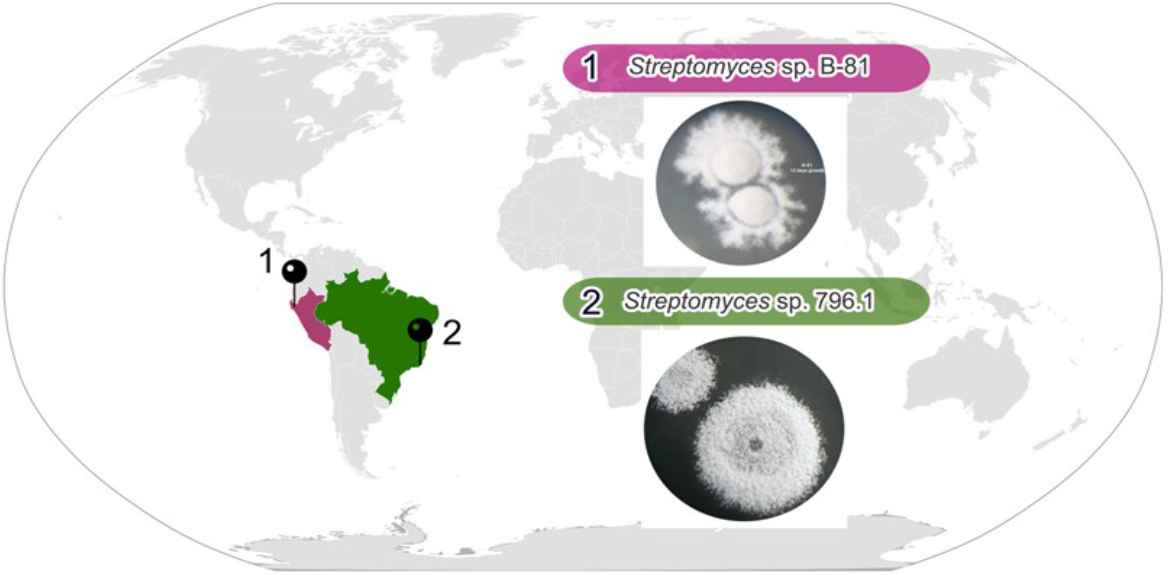
The geographical distribution of *Streptomyces* spp. location with a description of growth in solid medium. (1) Sediment of a Bayovar saline lagoon located in northwestern Peru and (2) starfish in Cabo Frio (RJ) in Brazil.

## 2. Results and Discussion

### 2.1. Phylogenetic analysis of 16S RNAr from two Streptomyces spp

The 16S rRNA sequences of the two marine *Streptomyces* species analyzed in this study, *Streptomyces* sp. B-81 collected in 2015 in Peru and *Streptomyces* sp. 796.1 collected in Brazil 2012, were determined and compared by MEGA11.0 software. The phylogenetic tree obtained from the two *Streptomyces* strains was mainly divided into two characteristic branches. The samples mentioned before were constructed against 29 *Streptomyces* strains according to EZBioCloud and are presented in Figure 2. The strain obtained from the Peruvian saline lagoon was determined to be *Streptomyces* sp. B-81 (MW562807), whose phylogenetically related sequence identity (1345 bp 92.7%, gene sequence identity) groups to the type strain *S. olivaceus* NRRL B-3009^T^ with 99,93%, *S. pactum* NBRC 13433^T^ and bootstrap value 97% (Fig. 2 and Table S1 and Table S2). However, for the strain isolated from a marine invertebrate in Brazil, *Streptomyces* sp. 796.1 (MG654686) sequence identity (1370 bp 93.4%, gene sequence identity similarity) is phylogenetically related to the type strains *S. buecherae* AC541^T^ with 99,70%, *S. youssoufiensis* X4^T^ and *S. zagrosensis* HM 1154^T^ with 99,48%, and *Streptomyces philanthi triangulum^T^* (DQ3752) with 99,45% gene sequence identity similarity and a bootstrap value of 85% (Figure 2 and Tables S1 and S2). Subsequently, each part was further subdivided, and most strains were well separated by clustering under each species, generating indications that they may be new species. Existing aquatic ecosystems are relatively unexplored but have been reported to harbor a great diversity of microorganisms, which may be reflected in their broad biosynthetic potential [17].

**Figure 2.**
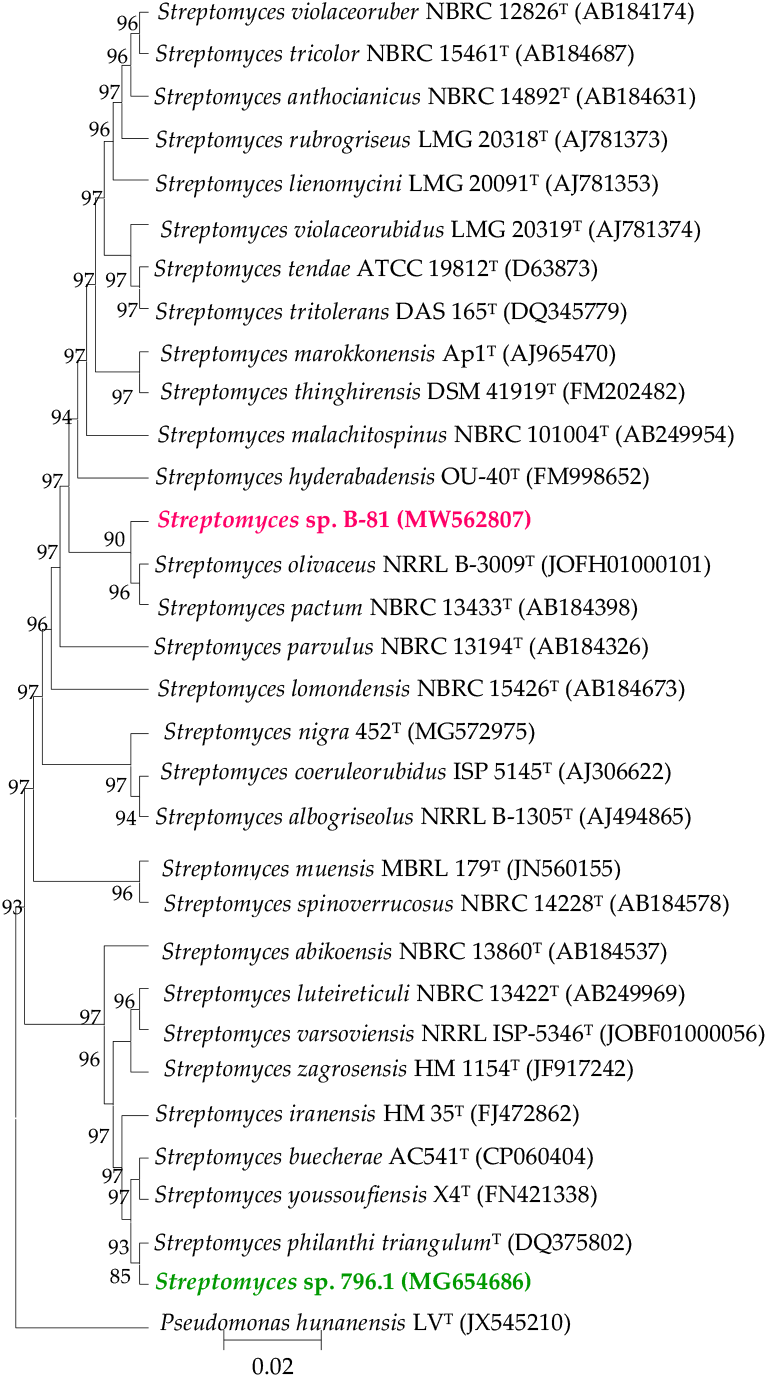
Phylogenetic tree of *Streptomyces* spp. strains B-81 and 796.1. The evolutionary history was inferred using the neighbor-joining method [18]. The percentage of replicate trees in which the associated taxa clustered together in the bootstrap test (1000 replicates) are shown next to the branches. The tree is drawn to scale, with branch lengths similar to evolutionary distances used to infer the phylogenetic tree. The evolutionary distances were computed using the Kimura 2-parameter method [19] and are in units of the number of base substitutions per site. The analysis involved 32 nucleotide sequences. All positions containing gaps and missing data were eliminated. There were a total of 1498 positions in the final dataset. Evolutionary analyses were conducted in MEGA11 [20]. Bar, 0.02 substitutions per nucleotide, *Pseudomonas hunanen-sis* LV^T^ (JX545210) was used as the outgroup.

### 2.2. MS/MS-Based Molecular Networks of Streptomyces spp

The metabolites of the crude extracts obtained from each of the *Streptomyces* spp. samples were prepared and analyzed by UHPLC–MS/MS, and the resulting data were analyzed through multivariate data analyses to compare both strains using LC– MS (Fig. 3). The data showed a clear separation between both *Streptomyces* spp. strains. In particular, using principal component analysis (PCA), *Streptomyces* sp. B-81 was separated from *Streptomyces* sp. 796.1 by PC1 and PC3, explaining ~ 40% of the variance (Fig. 3A). Another multivariate analysis, partial least squares-discriminant analysis (PLS-DA), showed clear differentiation between both *Streptomyces* strains using components 1 and 2 (Fig. 3B), and chemotaxonomic classification based on *Streptomyces* using liquid chromatography-electrospray ionization-tandem mass spectrometry (LC–ESI–MS/MS) combined with multivariate statistical analysis demonstrated that metabolite-based chemotaxonomic classification is an effective tool for distinguishing *Streptomyces* spp. and for determining their species-specific metabolites [21].

**Figure 3.**
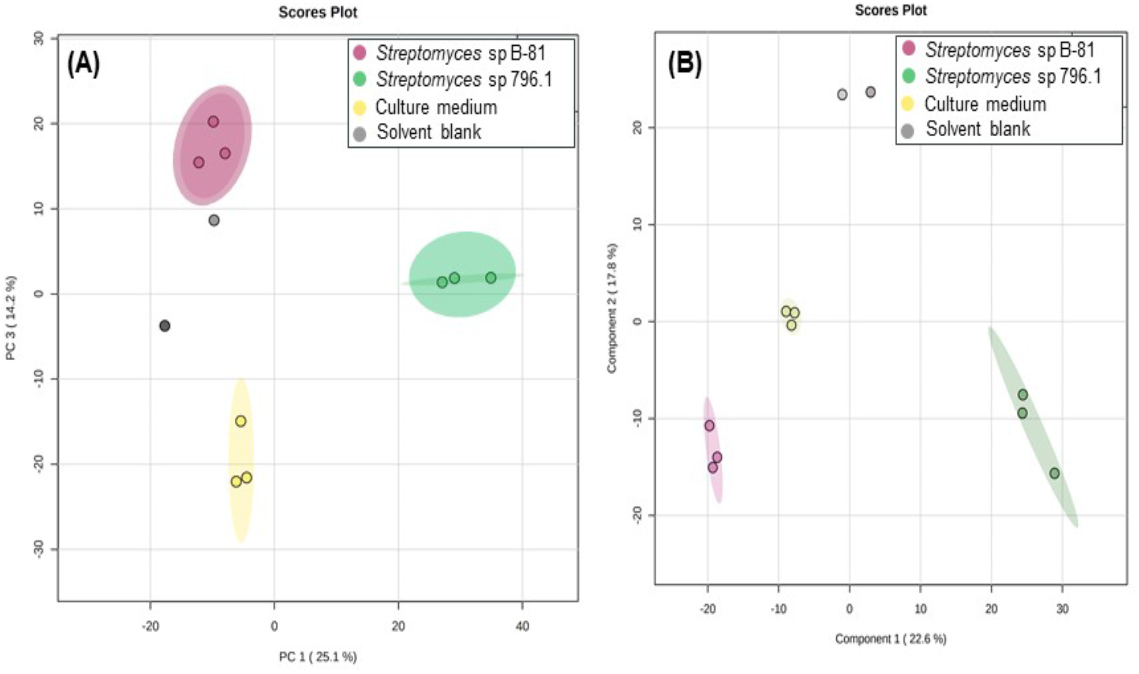
Multivariate data analysis of LC–MS/MS: (**A).** Principal component analysis (PCA) and (**B**). partial least square-discriminant analysis (PLS-DA) comparing both *Streptomyces* strains. *Streptomyces* sp. B81 is in pink, *Streptomyces* sp. 796.1, in green, the culture medium R2A, in yellow and the solvent blank is in white.

Mass spectrometric data obtained from these analyses were then used to generate molecular networking coupled to in silico tools (Networking annotation propagation (NAP) (https://ccms-ucsd.github.io/GNPSDocumentation/nap/) and MolNetEn-hancer (https://ccms-ucsd.github.io/GNPSDocumentation/molnetenhancer/). Data obtained from these analyses were then used to generate more detailed molecular networking (Fig. 4A). Our analyses have presented several chemical classes, such as lipids and lipid-like molecules, organic nitrogen organoheterocyclic compounds, organic acids, organosulfur compounds, polyketides, and benzenoid compounds (Fig. 4B).

**Figure 4.**
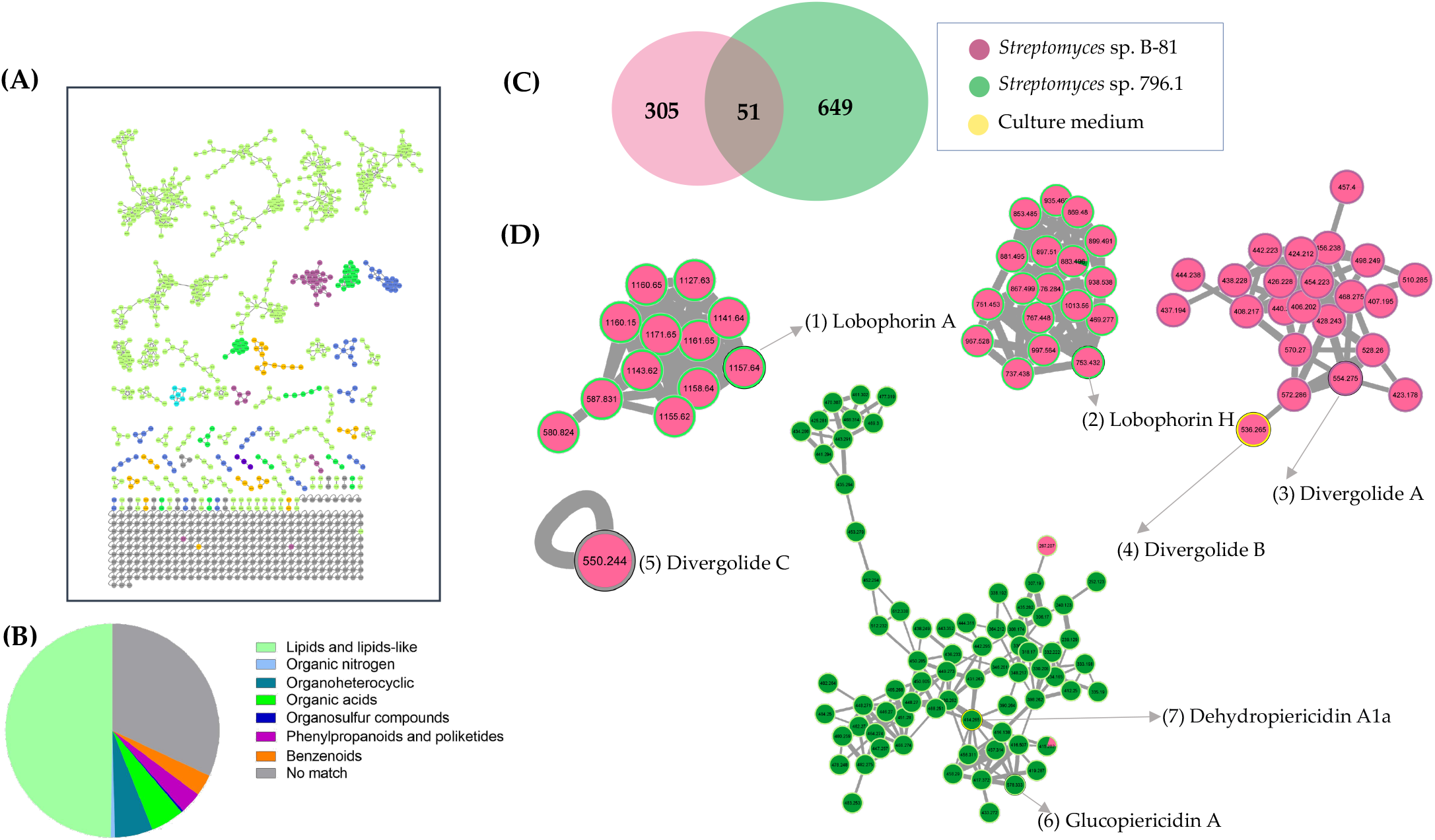
(**A**) MolNetEnhancer of *Streptomyces* strains LC–MS/MS data. (**B**) Pie chart based on the chemical classes found on the LC–MS/MS data. (**C**) Euler diagram based on the ion distribution of both strains. (**D**) Classical molecular networking of both *Streptomyces* strains.

Microorganisms in the environment can produce a wide range of secondary metabolites that are natural products with diverse chemical structures and that will perform a variety of functions acting as antibiotics, antitumour agents, cholesterol-lowering agents, etc. [22]. According to the LC–MS/MS analysis, the number of ions can be classified according to the strain source. In all 1005 detected ions, ~ 30% are only from *Streptomyces* sp. B-81 and ~ 64,5% from *Streptomyces* sp. 796.1 (Fig. 4C). Thus, compounds of the same chemical class are grouped in the same cluster, demonstrating a broad biomolecule profile for *Streptomyces* sp. B-81 (pink), including the production of lobophorins A (1) and H (2), annotated based on their accurate masses, with specific masses of 1157.6379 (1) and 753.4320 (2), respectively; metabolites were annotated as hits in the GNPS database or manually by accurate mass analyses, which showed mass errors below 0.10 ppm (Table 1). The cluster of the GNPS database indicated the production of lobophorin A by *Streptomyces* sp. B-81 (MW562807) strain, and fragmentation profile with typical fragments at 97.06, 183.11, 517.29 *m/z* (Fig. S1), the metabolite lobophorin H presented the precursor at *m/z* 753.4320 (Fig S2), and compounds that belong to the divergolide family, including divergolides A (3), B (4) and C (5), based on their precursor ions with *m/z* values of 554.2750 (3), 536.2650 (4), and 550.2440 (5), respectively. The fragmentation profile with typical fragments at divergolide C showed the main fragments *m*/*z* 182.08, 332.22, 398.32 (S3 – S5 Figs). In a previous study of Bayovar’s saloons *Streptomyces* sp. B-81 (MW562807) demonstrated the most promising activity, and six biomolecules (Cholic Acid, Lobophorin A, B, E, K and six compounds of type Furano) [14]. The MS/MS spectra of the network metabolites were compared with the GNPS spectral libraries and identified for *Streptomyces* sp. 796.1, two substances were annotated as glucopiericidin A (6) and dehydropiericidin A1a (7), with precursor ions at *m/z* 578.333 (7) and 414.26 (8), which were identified based on their fragmentation profiles with typical fragments at *m/z* 161.13, 182.09, 398.26 and 135.11, 182.09, 330.20, respectively (Figs S6–S7) (Fig. 4D and Table 1). In a study carried out on the marine sediment of Saint Peter and Saint Paul’s archipelago in Brazil, dereplication analysis of the extracts showed the presence of several compounds identified as piericidins A and C as well as glucopiericidin A, produced by *Streptomyces* sp. BRA-199 [23]

**Table 1.** Mass error (0,10 ppm) of observed and calculated *m/z* values of known compounds found in clusters based on MS/MS data obtained from UHPL-HRMS.

| N | GNPS NEW-ID | Compound | Molecular Formula [M+H]<sup>+</sup> | $m/z$ theoretical | $m/z$ observed | error (ppm) | Strain |
| --- | --- | --- | --- | --- | --- | --- | --- |
| 1 | 5653 | Lobophorin A | C61H93N2O19 | 1157.6367 | 1157.6379 | 1.032275 | B-81 |
| 2 | 5178 | Lobophorin H | C42H61N2O10 | 753.4320 | 753.43200 | 0.096890 | B-81 |
| 3 | 4097 | Divergolide A | C31H40NO8 | 554.2748 | 554.27500 | 0.281449 | B-81 |
| 4 | 3981 | Divergolide B | C31H38NO7 | 536.2642 | 536.26500 | 1.344486 | B-81 |
| 5 | 4068 | Divergolide C | C31H36NO8 | 550.2435 | 550.24400 | 0.828724 | B-81 |
| 6 | 4214 | Glucopiericidin A | C31H48NO9 | 578.3323 | 578.3330 | 1.108359 | 796.1 |
| 7 | 3145 | Dehydropiericidin A1a | C25H36NO4 | 414.263885 | 414.2650 | 2.691521 | 796.1 |

Error (0,10 ppm) = observed $m/z$ – calculated $m/z$ .

Ding and colleagues in 2018 took an approach combining LC–MS/MS and molecular networks as a rapid analytical method to detect six compounds. Aplysiatoxins from marine cyanobacteria were described with potential new analogs. Another study of MS/MS data obtained from fractions on the public GNPS platform succeeded in dereplicating new glycolipopeptides called characellides, cyanocobalamin and poecillastrins of a deep-sea tetractinellid sponge, *Characella pachastrelloides* [24,25]. Initial chemical investigation of the extract obtained from *Streptomyces* spp. indicated the production of compounds 1-7. Each node within the clusters represents an ionized metabolite, and dereplication was performed by comparison with the *m/z* values of known lobophorins, divergolides and piericidin-related analogs (Table 2). Lobophorins A and B were discovered by Jiang et al. (1999) [26] to Belize onboard the Columbus research ship. Lobophorin C and D were isolated from a new actinomycete strain named # CNC-837, identified as *Streptomyces carnosus* strain AZS17, isolated from the coastal waters of the East China Sea [27]. Two polyketides, spiro-tetronate classified as lobophorins H and I, were isolated from *Streptomyces* sp. 1053U.I.1a.3b [28], as well as the other substances of the Lobophorin family (Table 2). The family of compounds named Divergolides isolated from the stem of the mangrove tree *Streptomyces* sp. HKI0576 grown by HPLC–MS revealed a complex metabolome [8]. Four ansamycins, named divergolides T–W, isolated from the mangrove-derived actinomycete *Streptomyces* sp. KFD18 [29], a substance named 3’-deoxytalopiericidin A1 or glucopiericidin A, was isolated from *Streptomyces* sp. soil sample collected in Gotemba City, Shizuoka, Japan [30], 13-hydroxyglucopiericidin A and the analog Glucopiericidin A were isolated from the fermentation broth of *Streptomyces* sp. OM-5689 [31] (Table 2).

**Table 2.** A list of known compound families related to *Streptomyces* spp. with compound bioactivity with *m/z* values.

| Compound | Molecular Formula [M+H]<sup>+</sup> | Mass Molecular | Mass calculated | Origins | Reference |
| --- | --- | --- | --- | --- | --- |
| Lobophorin A | C61H93N2O19 | 1157.636705 | 1157.6379 | *Streptomyces* sp. CNC-837 | B-81, [26] |
| Lobophorin B | C61H91N2O21 | 1187.610884 | 1187.6114 | *Streptomyces* sp. CNC-837 | [26], [14] |
| Lobophorin C | C61H90N2O21 | 1186.603059 | 1209.5958 | *Streptomyces* sp. AZS 17 | [27] |
| Lobophorin D | C61H93N2O19 | 1157.636705 | 1157.6379 | *Streptomyces* sp. AZS 17 | [27] |
| Lobophorin E | C61H91N2O20 | 1171.615970 | 1171.6165 | *Streptomyces* sp. SCSIO 01127 | [32] |
| Lobophorin F | C54H78N2O17 | 1026.529500 | 1049.5198 | *Streptomyces* sp. SCSIO 01127 | [32] |
| Lobophorin G | C63H94N2O20 | 1198.639445 | 1199.6472 | *Streptomyces* sp. MS100061 | [33] |
| Lobophorin H | C42H61N2O10 | 753.432073 | 753.43200 | *Streptomyces* sp. 1053U.I.1a.3b | B-81, [28] |
| Lobophorin I | C61H89N2O21 | 1185.596209 | 1185.5958 | *Streptomyces* sp. 1053U.I.1a.3b | [28] |
| Lobophorin K | C61H93N2O20 | 1173.631620 | 1173.6322 | *Streptomyces* sp. M-207 | [34] |
| Lobophorin CR1 | C61H92NO20 | 1144.617647 | 1180.6032 | *Streptomyces* sp. 7790_N4 | [35] |
| Lobophorin CR2 | C61H90N2O22 | 1202.597974 | 1247.5702 | *Streptomyces* sp. 7790_N4 | [35] |
| Lobophorin CR3 | C61H90N2O23 | 1218.592889 | 1241.5832 | *Streptomyces* sp. 7790_N4 | [35] |
| Lobophorin CR4 | C52H77N2O15 | 969.531846 | - | *Streptomyces olivaceus* SCSIO T05 | [36] |
| Lobophorin L | C54H80N2O16 | 1013.5618 | 1013.5581 | *Streptomyces* sp. 4506 | [37] |
| Lobophorin M | C48H70N2O13 | 882.487242 | 883.4951 | *Streptomyces* sp. 4506 | [37] |
| Lobophorin N | C61H92N2O18 | 1141.64336 | 1141.6418 | *Streptomyces* sp. HDN1844000 | [38] |
| Divergolide A | C31H39NO8 | 554.274844 | 554.27500 | *Streptomyces* sp. HKI0576 | B-81, [8] |
| Divergolide B | C31H37NO7 | 536.264279 | 536.26500 | *Streptomyces* sp. HKI0576 | B-81, [8] |
| Divergolide C | C31H33NO8 | 550.243544 | 550.24400 | *Streptomyces* sp. HKI0576 | B-81, [8] |
| Divergolide D | C31H35NO8 | 535.232645 | 535.2326 | *Streptomyces* sp. HKI0576 | [8] |
| Divergolide T | C31H37NO7 | 536.264300 | 536.2641 | *Streptomyces* sp. KFD18 | [29] |
| Divergolide U | C31H37NO8 | 550.244600 | 550.2438 | *Streptomyces* sp. KFD18 | [29] |
| Divergolide V | C31H37NO7 | 536.264000 | 536.2640 | *Streptomyces* sp. KFD18 | [29] |
| Divergolide W | C31H37NO7 | 534.249000 | 534.2490 | *Streptomyces* sp. KFD18 | [29] |
| Piericidin A-A1 | C25H<sub>37-39</sub>NO4 | 401.268636 | 401.2686 | *Streptomyces* sp. 16-22 | [39] |
| Piericidin B-B1 | C26H39NO4 | 415.284286 | - | *Streptomyces mobaraensis* | [40] |
| Piericidins B1N-oxide | C26H39NO5 | 431.279201 | 446.2897 | *Streptomyces* sp. MJ288-OF3 | [41] |
| Piericidin C7 | C28H41NO5 | 457.294851 | 472.3062 | *Streptomyces* sp. YM14-060 | [42] |
| Piericidin C8 | C29H43NO5 | 471.310501 | 486.3223 | *Streptomyces* sp. YM14-060 | [42] |
| Glucopiericidin A | C31H48NO9 | 578.332359 | 578.3330 | *Streptomyces* spp. | 796.1, [30] |
| Dehydropiericidin A | C25H36NO4 | 414.263885 | 414.2650 | *Streptomyces* spp. | 796.1, [31] |
| 13-hydroxyglucopiericidin A | C31H47NO10 | 579.316374 | 593.3163 | *Streptomyces* sp. OM-5689 | [43] |
| Glucopiericidin C | C30H45NO8 | 533.310895 | 533.3108 | *Streptomyces* sp. B811 | [44] |
| JBIR-02 | C27H39NO3 | 411.289372 | 426.2979 | *Streptomyces* sp. ML55 | [45] |

## 3. Materials and Methods

### 3.1. Biological Material

The microorganisms used in this study were preserved by deep freezing at −80°C in 30% glycerol in the research collection of the Microbial Resources Division, Pluridisciplinary Center for Chemical, Biological and Agricultural Research (CPQBA) State University of Campinas (UNICAMP), Campinas, SP, Brazil. (https://www.cpqba.unicamp.br/) and collection of CIICAM Research Center (www.ciicam.com). Two bacteria belonging to the genus *Streptomyces* were identified as *Streptomyces* sp. 796.1 (MG654686), isolated from a starfish in Cabo Frio (RJ) in Brazil in 2012, *Streptomyces* sp. B-81 (MW562807) was isolated from the sediment of a Bayovar saline lagoon located in northwestern Peru in 2015.

The isolates were reactivated using R2A medium (KASVI) with artificial sea-water (ASW) (MgCl_2_-6H_2_O 10.83 g. L^-1^, CaCl_2_-2H_2_O 1.51 g. L-^1^, SrCl2-6H_2_O 0.02 g. L-^1^, NaCl 23.93 g. L-^1^, Na_2_SO_4_ 4.01 g. L-^1^, KCl 0.68 g. L-^1^, NaHCO_3_ 0.20 g. L-^1^, KBr 0.098 g. L^-^1, H_3_BO_3_ 0.03 g. L^-1^) with 5% NaCl and incubated at 28°C for 15 days. After incubation, the colonies verified for purity were collected from the plate and used for the extraction of genomic DNA.

### 3.2. DNA isolation and PCR amplification of bacteria

The extraction of genomic DNA was performed using the protocol established by Van Soolingen et al. (1991) adapted to the conditions of the laboratory (https://dx.doi.org/10.17504/protocols.io.bsj4ncqw). First, quantification of genomic DNA was performed by comparing the Λ DNA at different concentrations in a 1% TBE 1X agarose gel submitted to electrophoresis for 20 min at 5 V per cm^2^. Then, the DNA was kept at −20°C until the amplification of the 16S rRNA gene was performed by polymerase chain reaction (PCR). The specifications of this protocol have been previously described (dx.doi.org/10.17504/protocols.io.brrmm546).

### 3.3. Sequencing and Phylogenetic Analysis of 16

The samples were purified using a minicolumns GFX PCR DNA & gel band purification kit (GE Healthcare Bio-Sciences AB Uppsala, Sweden) according to a previously described protocol (https://dx.doi.org/10.17504/protocols.io.brzpm75n). The samples were sequenced using an ABI3500XL Series automatic sequencer (Applied Biosystems Foster City, California, USA) in the Laboratory of the Division of Microbial Resources, Chemical, Biological and Agricultural Pluridisciplinary Research Center (CPQBA), University of Campinas (UNICAMP), Paulínia, São Paulo, Brazil. Sequencing reactions were performed using the Big Dye Terminator Cycle Sequencing Ready Reaction Kit (Applied Biosystems) according to the manufacturer’s modified by the author’s protocol (https://dx.doi.org/10.17504/protocols.io.brzpm75n).

Partial sequences of the 16S ribosomal RNA gene obtained from each isolate were assembled into a contig using BioEdit 7.0. The sequences of organisms were added to the EZBioCloud 16S Database (https://www.ezbiocloud.net/) using the “Identify” service, and species assignment was based on the closest hit 16S rRNA gene sequences retrieved from the database and related to the unknown organism gene were selected for alignment in the Clustal W program. Phylogenetic analyses were performed using the MEGA version 11.0 program, and the evolutionary distance matrix was calculated using the Kimura-2 model parameters. Then, the phylogenetic tree was constructed from the evolutionary distances calculated by the neighbor-joining method with bootstrap values based on 1000 resamples.

### 3.4. Production of Crude Extracts of Streptomyces spp

The production of crude ethyl acetate extracts of bacterial isolates was performed in culture media R2A broth (Himedia ref. 1687) for *Streptomyces* spp. B-81 and 796.1 strains as described in a previously published protocol (dx.doi.org/10.17504/protocols.io.br2cm8aw). *Streptomyces* sp. metabolites were extracted with 1000 μL of methanol in an ultrasonic bath for 40 minutes, dried under inert conditions and analyzed by ultrahigh pressure liquid chromatography–mass spectrometry (UHPLC-MS) in a Thermo Scientific QExactive^®^ Hybrid Quadrupole-Orbitrap Mass Spectrometer.

### 3.5. LC–MS/MS data acquisition

Analyses were performed in the positive mode with a range of m/z 133-2000, a capillary voltage of 3.4 kV, inlet capillary temperature of 280°C, and S-lens 100 V. A Thermo Scientific column Accucore C18 2.6 μm (2.1 mm x 100 mm) was used as the stationary phase, and 5 μL of the sample was injected. The mobile phase was composed of 0.1% formic acid and acetonitrile, and the eluent profile followed a molar ratio of 95/5 up to 2/98 within 10 min, held for 5 min, up to 95/5 within 1.2 min, and held for 8.8 min. The total run time was 25 min for each run, and the flow rate was 0.2 mL x min-1. The data were initially processed with Xcalibur software (version 3.0.63) developed by Thermo Fisher Scientific.

### 3.6. LC–MS/MS data analysis and metabolite annotation

Raw data, blanks, and media controls were converted to.mzML using MSCon-vert software (http://proteowizard.sourceforge.net). Next, a molecular network for *Streptomyces* sp. strain metabolites was created using the online workflow (https://ccms-ucsd.github.io/GNPSDocumentation/networking/) on the Global Natural Products Social Molecular Networking (GNPS) website platform (http://gnps.ucsd.edu) using the Classical Molecular Networking (CMN) tool. For network creation, the data were filtered by removing all MS/MS fragment ions within +/− 17 Da of the precursor m/z. MS/MS spectra were filtered by choosing only the top 6 fragment ions in the +/− 50 Da window throughout the spectrum. The precursor ion mass tolerance was set to 0.02 Da and an MS/MS fragment ion tolerance of 0.02 Da. A network was then created where edges were filtered with a cosine score above 0.65 and more than six (6) matched peaks.

Furthermore, edges between two nodes were kept in the network if and only if each of the nodes appeared in each other’s respective top 10 most similar nodes. Finally, the maximum size of a molecular family was set to 100, and the lowest-scoring edges were removed from molecular families until the molecular family size was below this threshold. The spectra in the network were then searched against GNPS’ spectral libraries. The library spectra were filtered in the same manner as the input data. All matches kept between network spectra and library spectra were required to have a score above 0.65 and at least six (6) matched peaks [46]. The resulting molecular networking is available at:

(https://gnps.ucsd.edu/ProteoSAFe/status.jsp?task=b7a29b5a78864223bde01fb8af0c5eff). To perform the multivariate data analysis, the LC–MS data were processed into MZmine online using GNPS Dash-board (https://ccms-ucsd.github.io/GNPSDocumentation/lcms-dashboard/). The job is available at:

https://gnps.ucsd.edu/ProteoSAFe/status.jsp?task=88a21c1aabb04a1287348300b711ebfd. The quantification table was submitted to the MetaboAnalyst 5.0 platform (https://www.metaboanalyst.ca/) for statistical analysis.

The nodes (MS/MS spectra) originating from R2A culture media and blanks (methanol) were filtered from the original network to enable visualization of metabolites. Finally, the final spectral network (.cys) was uploaded in Cytoscape 3.8 to obtain better visualization and editing.

## 4. Conclusions

In this study, we sought to classify two *Streptomyces* species based on their secondary metabolites. The dendrograms based on metabolites and 16S rRNA exhibited the same patterns of species discrimination and may be new species. The multivariate data analyses to compare both strains using LC–MS showed a clear separation between both *Streptomyces* spp. The MS/MS spectra of the network metabolites were compared with the GNPS spectral libraries, and differences in the molecular network presented in *Streptomyces* sp. B-81 isolated from the Bayovar saline lagoon located in northwestern Peru were lobophorins A (1) and H (2), as well as divergolides A (3), B (4) and C (5); *Streptomyces* sp. 796.1 isolated in Cabo Frio (RJ) in Brazil dereplicated two compounds, glucopiericidin A (6) and dehydropiericidin A1a (7).

## Supplementary Materials

The following supporting information can be downloaded at www.mdpi.com/xxx/s1. Phylogenetic data of *Streptomyces* sp. B-81 Accession number (MW562807), and *Streptomyces* sp. 796.1 Accession number (MG654686); MS/MS match between GNPS database lobophorin A (1), in the GNPS database or manually by accurate determination of which ion precursor lobophorin H (2) and ions precursor divergolides A (3), B (4) and C (5) from *Streptomyces* sp. B-81 extract and MS/MS match between GNPS database with Glucopiericidin A (6) and Dehydropiericidin A1a (7).

## Author Contributions

R.F.C. performed the extraction of *Streptomyces* sp. B-81 sample, the isolation, purification, and structure elucidation of compounds, and constructed the molecular networks A.K.P, wrote and paper; N.R.Q. performed the isolation and production of *Streptomyces* sp. 796.1 extracts. A.K.P. and J.H.C performed the UHPLC–MS/MS analysis and Thermo Scientific QExactive^®^ Hybrid Quadrupole-Orbitrap Mass Spectrometer and constructed the molecular networks; T.P.F and F.F.G edited and vetted the paper. All authors read and corrected the paper.

## Acknowledgments

This work was supported by the Biotechnology Department of the Center for Research and Innovation in Multidisciplinary Active Sciences -CIICAM and Pluridisciplinary Center for Chemical, Biological and Agricultural Research (CPQBA), State University of Campinas (UNICAMP), Paulínia, SP, Brazil.

## SUPPLEMENTARY DATA

**Table S1.**
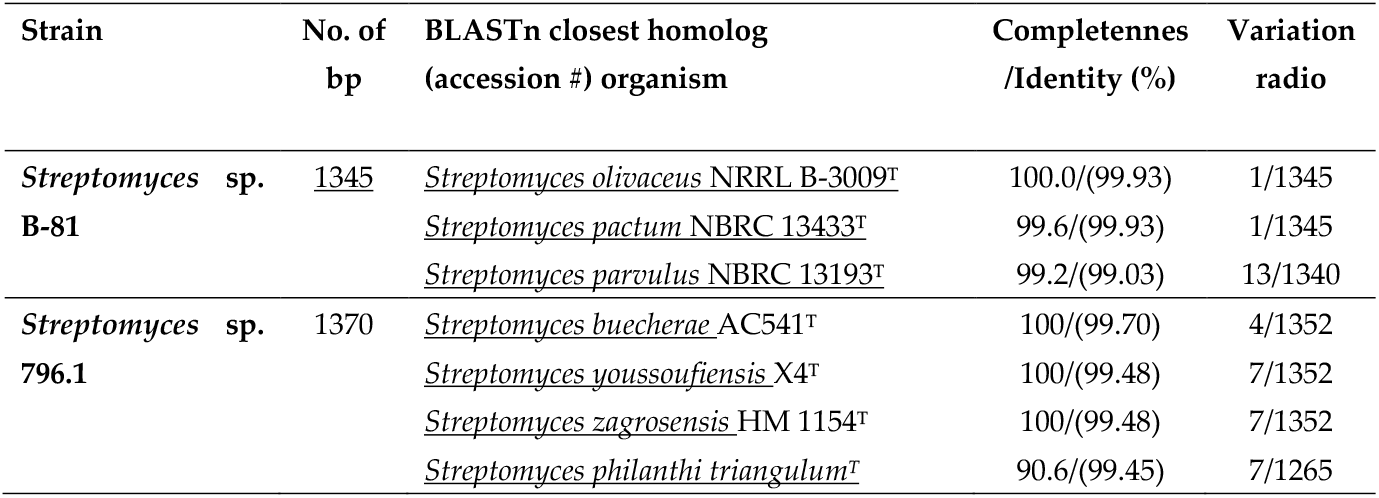
Complete 16S rRNA sequences of two *Streptomyces* spp.

**Table S2.**
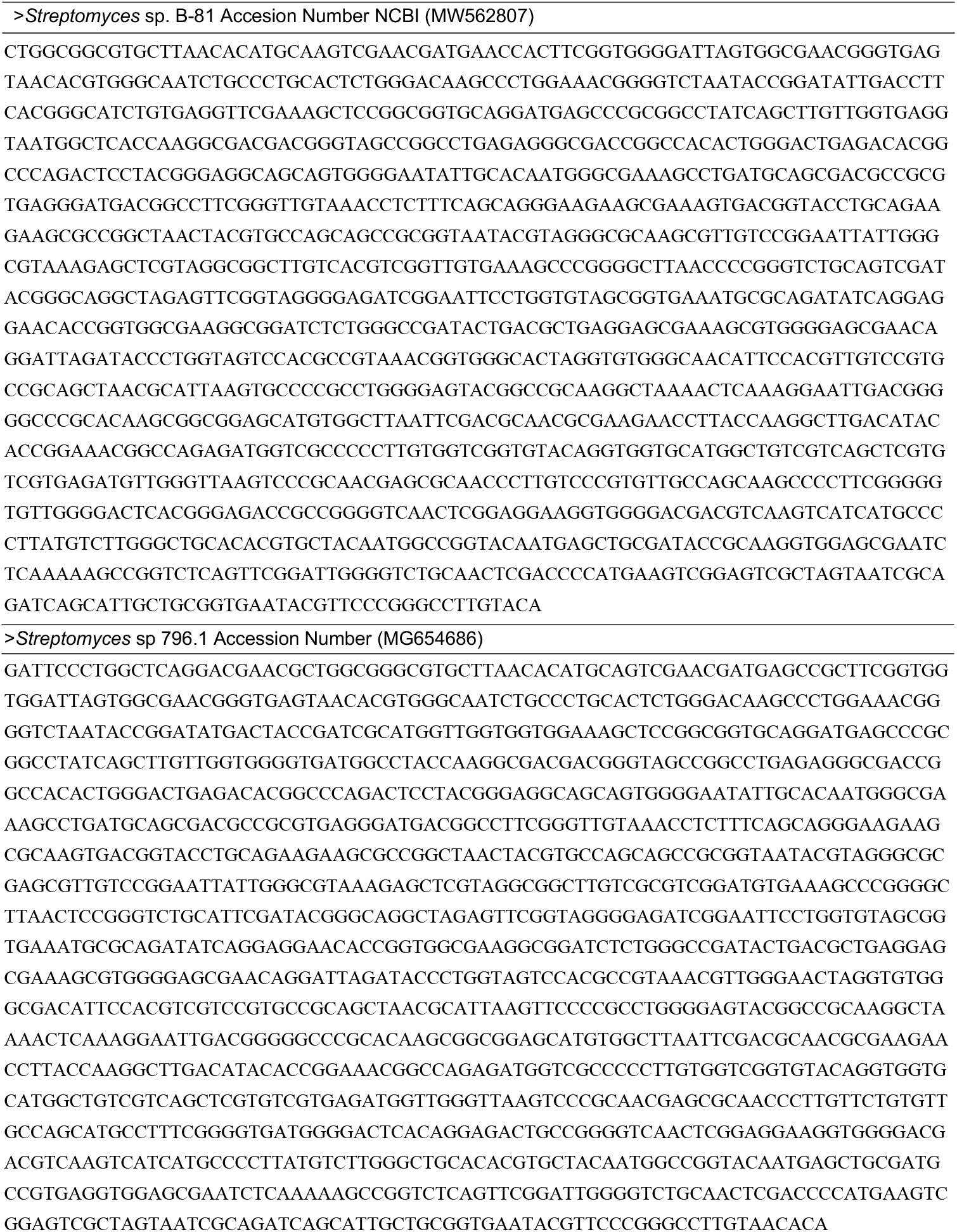
Complete 16S rRNA sequences of three *Streptomyces* spp.

**Fig. S1.**
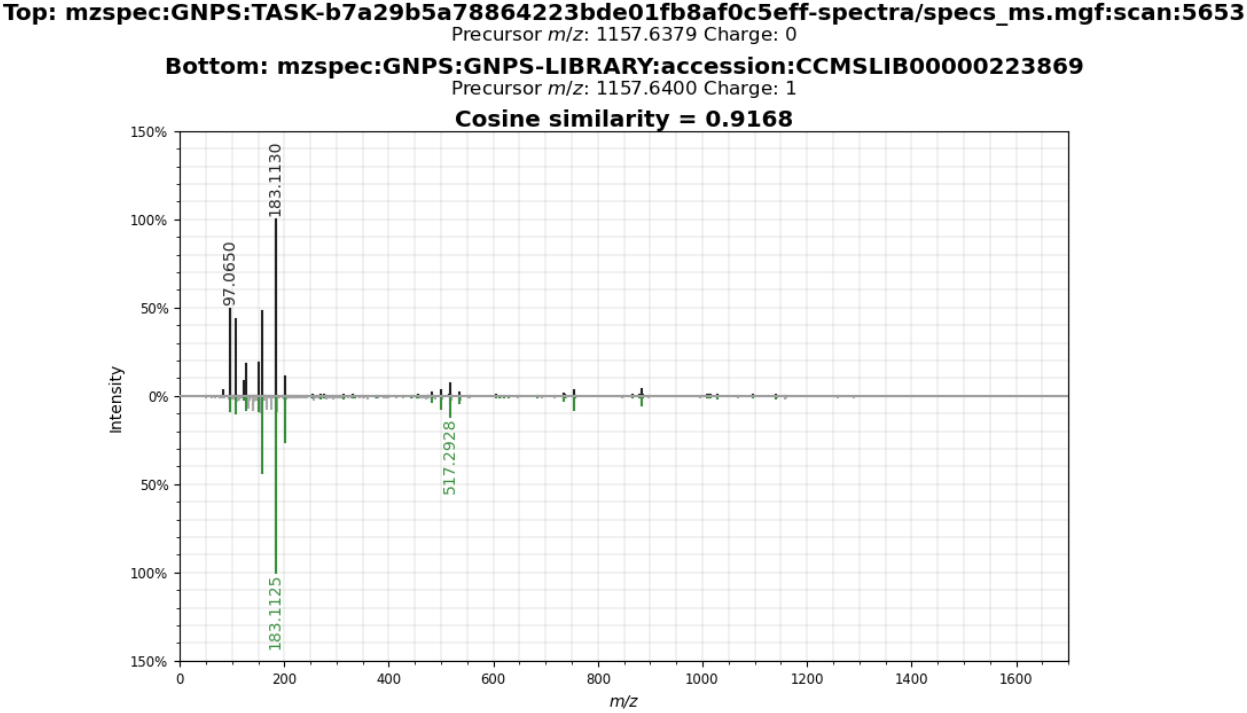
MS/MS match between GNPS database (green) lobophorin A C61H93N2O19 (5653) compound (1) from *Streptomyces* sp. B-81 extract (black)

**Fig. S2.**
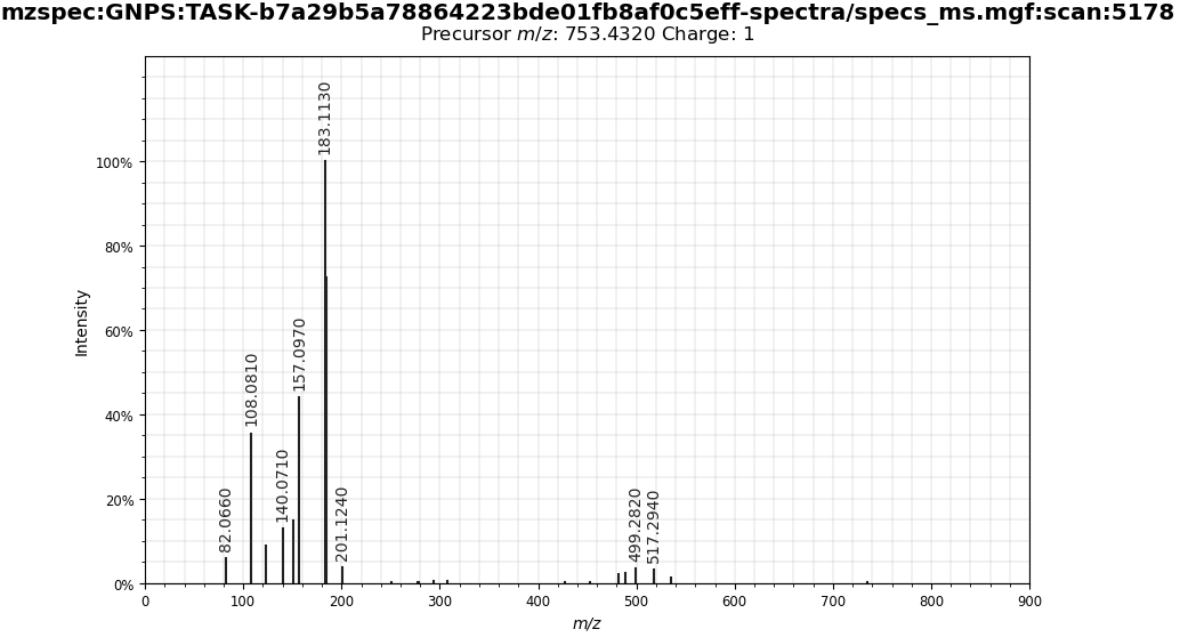
Lobophorin H in the GNPS database or manually by accurately determining which ion precursor m/z 753.4320 showed mass errors below 0.10 ppm C42H60N2O10 (5178) from *Streptomyces* sp. B-81 extract.

**Fig. S3.**
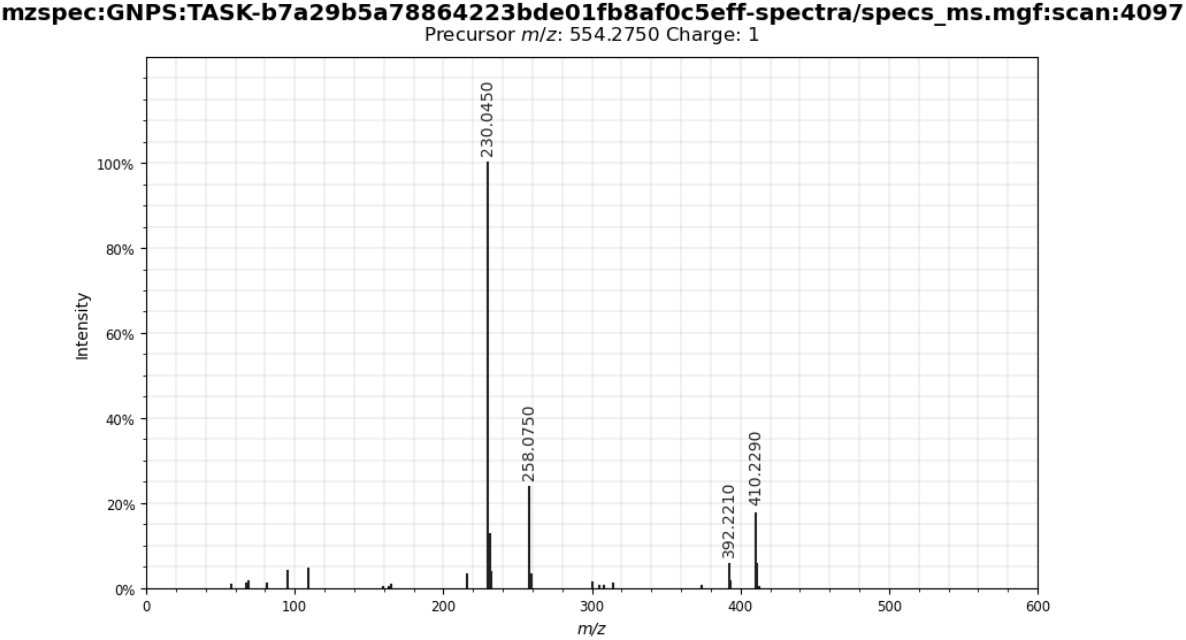
Divergolide A in the GNPS database or manually by accurate determination of which ion precursor m/z 554.2750 showed mass errors below 0.10 ppm C31H39NO8 (4097) compound (3) from *Streptomyces* sp. B-81 extract.

**Fig. S4.**
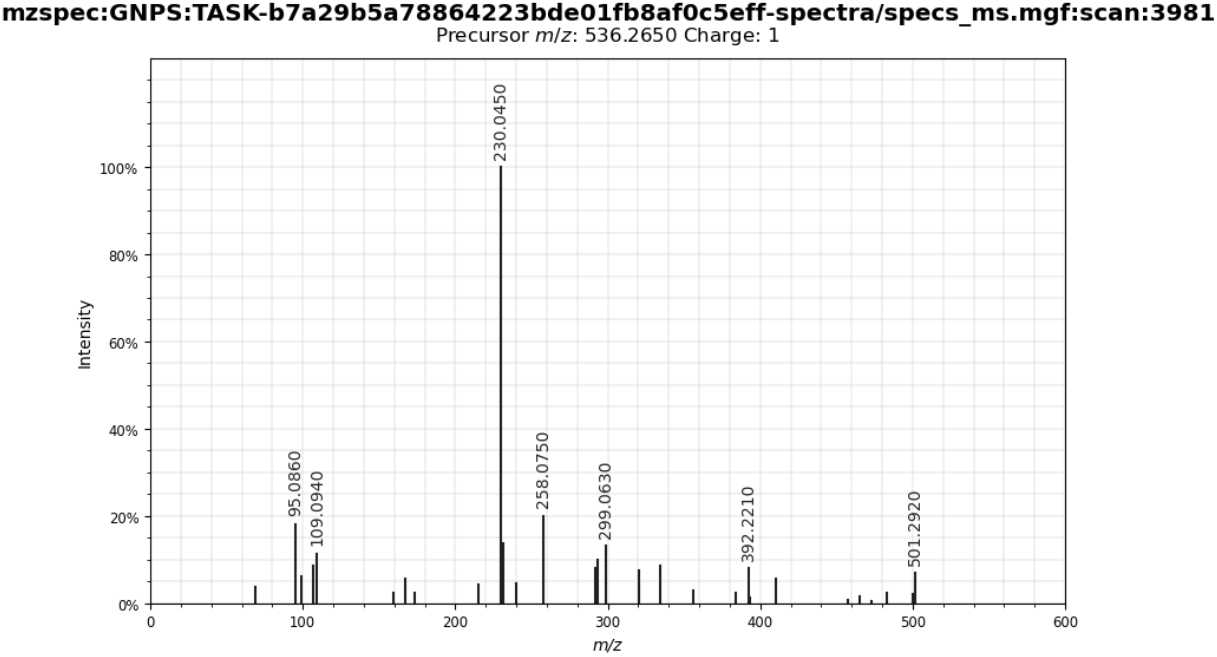
Divergolide B in the GNPS database or manually by accurate determination of which ion precursor m/z 536.2650 showed mass errors below 0.10 ppm C31H37NO7 (3981) compound (4) from Streptomyces sp. B-81 extract.

**Fig. S5.**
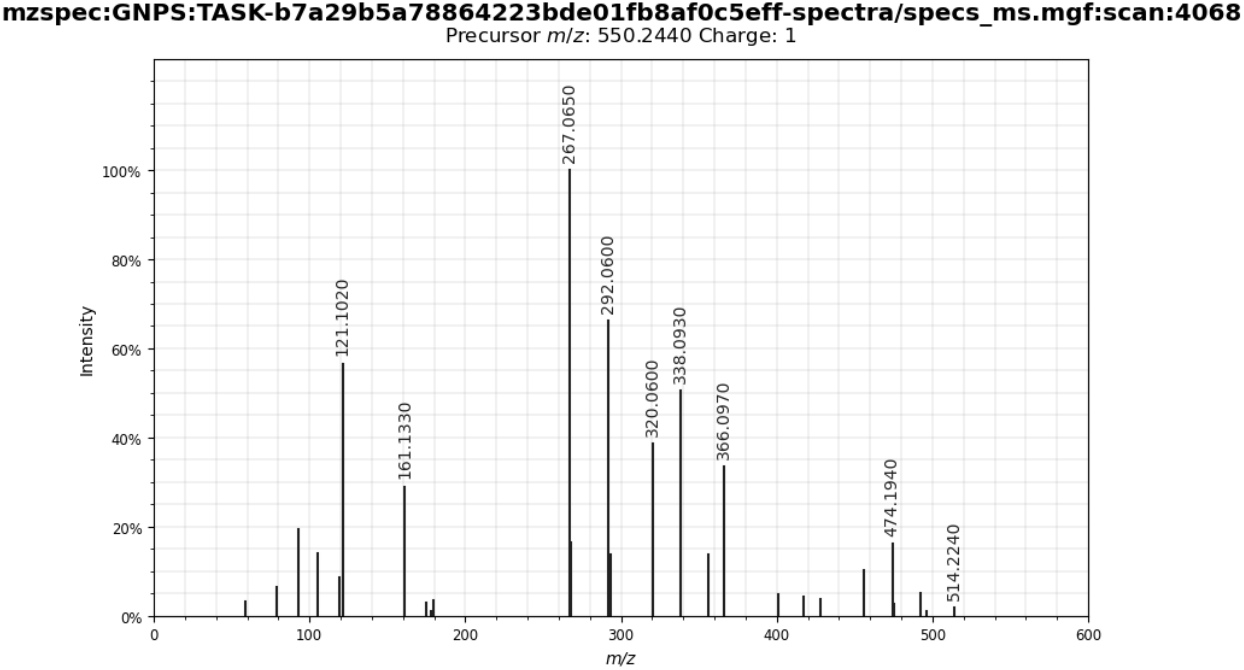
Divergolide C in the GNPS database or manually by accurate determination of which ion precursor m/z 550.2440 showed mass errors below 0.10 ppm C31H35NO8 (4068) compound (5) from *Streptomyces* sp. B-81 extract.

**Fig. S6.**
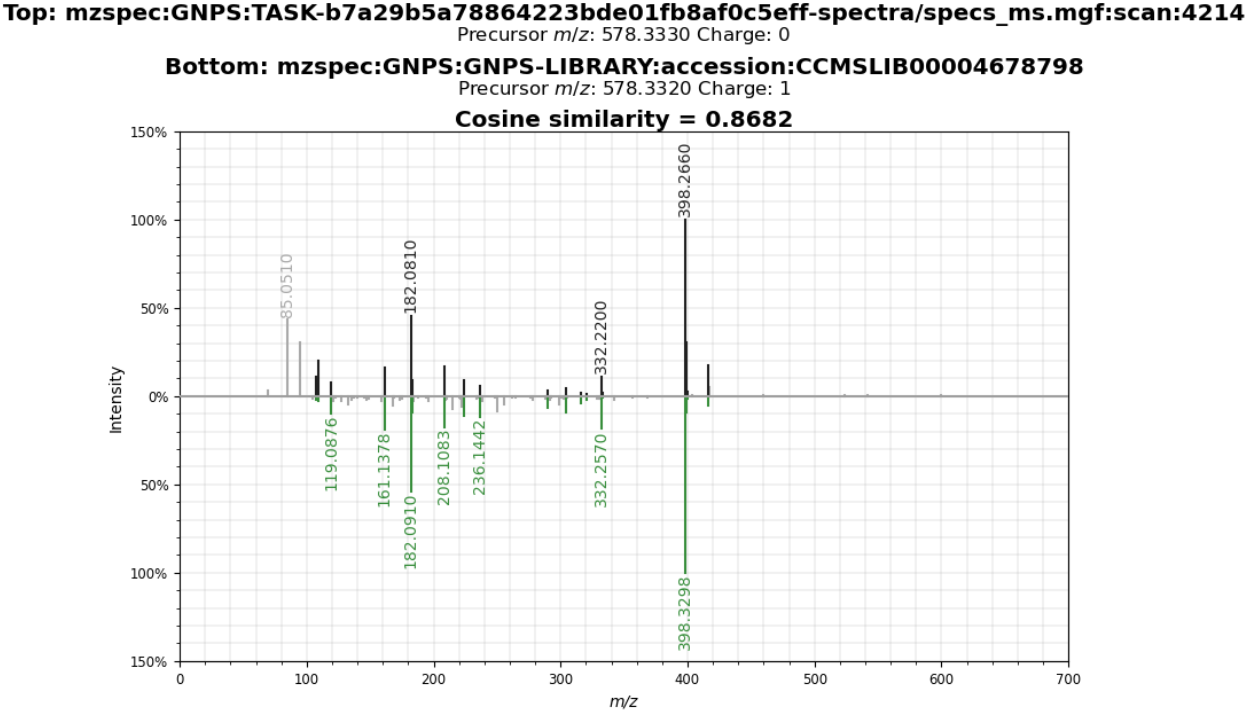
MS/MS match between GNPS database (green) glucopiericidin A C31H47NO9 (4214) compound (6) from *Streptomyces* sp. 796.1 extract (black)

**Fig. S7.**
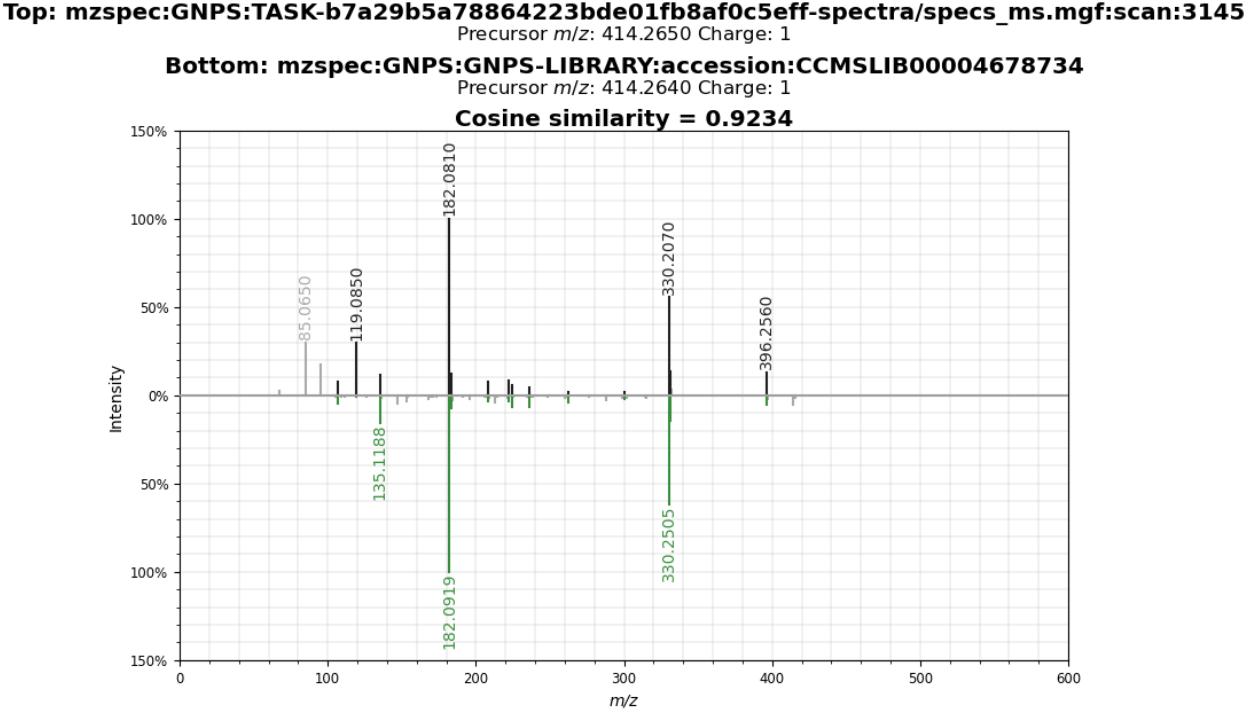
MS/MS match between GNPS database (green) dehydropiericidin A1a C25H35NO4 (3145) compound (**7**) from *Streptomyces* sp. 796.1 extract (black).

## References

1. Hayakawa, Y; Shirasaki, S; Shiba, S; Kawasaki, T; Matsuo, Y; Adachi, K., & Sishuri, Y. Piericidins C7 and C8, new cytotoxic antibiotics produced by a marine *Streptomyces* sp. J. Antibiot, 2007, 60, 196–200. doi:10.1038/ja.2007.22. [PubMed]

2. Valliappan, K; Sun, W; Li, Z. Marine actinobacteria associated with marine organisms and their potentials in producing pharmaceutical natural products. Appl. Microbiol. Biotechnol. 2014, 98, 7365–7377. doi:10.1007/s00253-014-5954-6. [PubMed]

3. Binayke, A; Ghorbel, S; Hmidet, N; Raut, A; Gunjal, A; Uzgare, A.,… & Nawani, N. Analysis of diversity of actinomycetes from arid and saline soils at Rajasthan, India. Environ. Sustain. 2018, 1–10. doi:10.1007/s42398-018-0003-5. [Crossref]

4. Kiyoon Kim; Sandipan Samaddar; Poulami Chatterjeea; Ramasamy Krishnamoorthy; Sunyoung Jeon, T.S. Structural and functional responses of microbial community with respect to salinity levels in a coastal reclamation land. Appl. Soil. Ecol. 2019, 137, 96–105. doi:10.1016/j.apsoil.2019.02.011. [Crossref]

5. Berdi, J. Bioactive microbial metabolites. Journal of antibiotics. 2015, 58, 1, 1–26. doi:10.1038/ja.2005.1. [Crossref] [PubMed]

6. Kamjam, M; Sivalingam, P; Deng, Z; Hong, K. Deep sea actinomycetes and their secondary metabolites. Front. Microbiol. 2017, 8, 1–9. doi:10.3389/fmicb.2017.00760. [Crossref] [PubMed]

7. Tangerina, M.M.P, Furtado, L.C, Leite, V.M.B, Bauermeister, A; Velasco-Alzate, K; Jimenez, P.C.… & Pena Ferreira, M.J. Metabolomic study of marine *Streptomyces* sp.: Secondary metabolites and the production of potential anticancer compounds. PLoS One. 2021, 15, 1–19. doi:10.1371/journal.pone.0244385. [Crossref] [PubMed]

8. Ding, L; Maier, A; Fiebig, H.H; Görls, H; Lin, W.H; Peschel, G., & Hertweck, C. Divergolides A-D from a mangrove endophyte reveal an unparalleled plasticity in ansa-macrolide biosynthesis. Angewandte Chemie. 2011, 50, 1630–1634. doi:10.1002/anie.201006165. [Crossref] [PubMed]

9. Hua-Qi, Pan; Song-Ya, Zhang; Nan, Wang; Zhan-Lin Li; Hui-Ming; Hua J-CH and Wang, S.J. “New spirotetronate antibiotics, lobophorins H and I, from a South China Sea-derived *Streptomyces* sp. 12A35. Marine Drugs, 2013. 3891–3901. doi:10.3390/md11103891. [Crossref] [PubMed]

10. Masaru Matsumoto; Kin-Ichi Mogi; Katsuhiko Nagaoka; Seiji Ishizeki, R.K and T.N. New Piericidin Glucosides, Glucopiericidins A and B. J. Antibiot. 1987, 149–156. doi:10.7164/antibiotics.40.149. [Crossref] [PubMed]

11. Braña, A.F; Sarmiento-Vizcaíno, A; Osset, M; Pérez-Victoria, I; Martín, J; De Pedro, N.… & Blanco, G. Lobophorin K, a new natural product with cytotoxic activity produced by *Streptomyces* sp. M-207 associated with the deep-sea coral Lophelia pertusa. Marine Drugs. 2017. doi:10.3390/md15050144. [Crossref] [PubMed]

12. Zhang, C; Ding, W; Qin, X; Ju, J. Genome Sequencing of *Streptomyces olivaceus* SCSIO T05 and Activated Production of Lobophorin CR4 via Metabolic Engineering and Genome Mining. Mar. Drugs. 2019, 17, 1–11. doi:10.3390/md17100593. [Crossref] [PubMed]

13. Carlos, Cortés-Albayay; Silver, J; Imhoff, J.F; Asenjo, J.A; Andrews, B; Nouioui, I., & Dorador, C. The Polyextreme Ecosystem, Salar de Huasco at the Chilean Altiplano of the Atacama Desert Houses Diverse *Streptomyces* spp. with Promising Pharmaceutical Potentials. Diversity. 2019, 11, 1–12. doi:10.3390/d11050069. [Crossref]

14. Flores Clavo, R; Ruiz Quiñones, N; Hernández-Tasco, A.J; Jose Salvador, M; Tasca Gois Ruiz, A.L; A.L, De Oliveira Braga; L.E.… & Fantinatti Garboggnini, F. Evaluation of antimicrobial and antiproliferative activities of Actinobacteria isolated from the saline lagoons of northwestern Peru. PLoS One. 2021. 1–18. doi: https://doi.org/10.1371/journal.pone.0240946. [Crossref] [PubMed]

15. Lee, S.Y; Kim, H.U; Chae, T.U; Cho, J.S; Kim, J.W; Shin, J.H.… & Jang, Y.S. A comprehensive metabolic map for production of biobased chemicals. Nat. Catal. 2019, 2, 18–33. doi:10.1038/s41929-018-0212-4. [Crossref]

16. Du Chao. Multiomics studies of the control of growth and antibiotic production of Streptomyces. (Doctoral dissertation Leiden University). 2020, 1–176. [Crossref]

17. Cuadrat, R.R.C; Ionescu, D; Dávila, A.M.R; Grossart, H.P. Recovering genomics clusters of secondary metabolites from lakes using genome-resolved metagenomics. Front. Microbiol. 2018, 9, 1–13. doi:10.3389/fmicb.2018.00251. [Crossref] [PubMed]

18. Saitou, N; Nei, M. The neighbor-joining method: a new method for reconstructing phylogenetic trees. Mol. Biol. Evol. 1987, 4, 406–425. doi:10.1093/oxfordjournals.molbev.a040454. [Crossref] [PubMed]

19. Kimura, M. A simple method for estimating evolutionary rates of base substitutions through comparative studies of nucleotide sequences. J. Mol Evol. 1980, 16, 111–120. doi:10.1007/BF01731581. [Crossref] [PubMed]

20. Tamura K; Stecher, G; Kumar, S. MEGA11: Molecular Evolutionary Genetics Analysis Version 11. Mol. Biol. Evol. 2021, 38, 3022–3027. doi:10.1093/molbev/msab120. [Crossref] [PubMed]

21. Lee, M.Y; Kim, H.Y; Lee, S; Kim, J.G; Suh, J.W; Lee, C.H. Metabolomics-based chemotaxonomic classification of *Streptomyces* spp. and its correlation with antibacterial activity. J. Microbiol. Biotechnol. 2015, 25, 1265–1274. doi:10.4014/jmb.1503.03005. [Crossref] [PubMed]

22. Chen, R; Wong, H; Burns, B. New Approaches to Detect Biosynthetic Gene Clusters in the Environment. Medicines. 2019, 6, 32. doi:10.3390/medicines6010032. [Crossref] [PubMed]

23. Ferreira, E.G; Torres, M da C.M; da Silva A.B; Colares, L.L.F; Pires, K; Lotufo, T.M.C.… & Jimenez, P.C. Prospecting Anti-cancer Compounds in Actinomycetes Recovered from the Sediments of Saint Peter and Saint Paul’s Archipelago, Brazil. Chem Biodivers. 2016, 13, 1149–1157. doi:10.1002/cbdv.201500514. [Crossref] [PubMed]

24. Ding, C.Y.G; Pang, L.M; Liang, Z.X; Goh, K.K.K; Glukhov, E; Gerwick, W.H; Tan, T.L. MS/MS-Based molecular networking approach for the detection of aplysiatoxin-Related compounds in environmental marine cyanobacteria. Mar Drugs. 2018,16, 1–15. doi:10.3390/md16120505. [Crossref] [PubMed]

25. Afoullouss, S; Sanchez, A.R; Jennings, L.K; Kee, Y; Allcock, A.L; Thomas O.P. Unveiling the Chemical Diversity of the Deep-Sea Sponge Characella pachastrelloides. Marine Drugs. 2022, 20, 1–52. doi:10.3390/md20010052. [Crossref] [PubMed]

26. Zhi-Dong Jiang, Paul R. Jensen and WF. Lobophorins A and B, new antiinflammatory macrolides produced by a tropical marine bacterium. Bioorganic & Medicinal Chemistry Letters. 1999, 9 (14), 2003–2006. [Crossref] [PubMed]

27. Rong-Bian Wei, Tao Xi, Jing Li, Ping Wang, Fu-Chao Li Y-CL and Qin, S. Lobophorin C and D, New Kijanimicin Derivatives from a Marine Sponge-Associated Actinomycetal Strain AZS17. Marine Drugs. 2011, 9 (3), 359–368. doi:10.3390/md9030359. [Crossref] [PubMed]

28. Lin, Z; Koch, M; Pond, C.D; Mabeza, G; Seronay, R.A; Concepcion, G.P.… & Schmidt, E.W. Structure and activity of lobophorins from a turrid mollusk-associated *Streptomyces* sp. Journal of Antibiotics. 2014, 67(1), 121–126. doi:10.1038/ja.2013.115. [Crossref] [PubMed]

29. Li-Man Zhou, Fan-Dong Kong, Qing-Yi Xie, Qing-Yun Ma, Zhong Hu, You-Xing Zhao and Luo, D.Q. Divergolides T–W with Apoptosis Inducing Activity from the Mangrove Derived Actinomycete *Streptomyces* sp. KFD18. Marine Drugs. 2019, 17(4), 1–9. doi:10.3390/md17040219. [Crossref] [PubMed]

30. Iwasaki, H; Kamisango, K.I; Kuboniwa, H; Sasaki, H; Matsubara, S. 3’-Deoxytalopiericidin a1 a novel analog of antitumor antibiotics from oligotroph. J. Antibiot. 1991, 44, 451–452. doi:10.7164/antibiotics.44.451. [Crossref] [PubMed]

31. Iroharu Mori; Shinji Funayama YS; Omura, K.K and Omura, S. A new antibiotic, 13-Hydroxyglucopiericidin A isolation, structure, elucidation and biological characteristics. J. Antibiot. 1990, XLIII, 1329–1331. [Crossref] [PubMed]

32. Niu, S; Li, S; Chen, Y; Tian, X; Zhang, H; Zhang, G.,… & Zhang, C. Lobophorins E and F, new spirotetronate antibiotics from a South China Sea-derived *Streptomyces* sp. SCSIO 01127. J. Antibiot. 2011, 64(11), 711–716. doi:10.1038/ja.2011.78. [Crossref] [PubMed]

33. Chen, C; Wang, J; Guo, H; Hou; Yang, N; Ren, B.… & Zhang, L. Three antimycobacterial metabolites identified from a marine-derived *Streptomyces* sp. MS100061. Applied microbiology and biotechnology. 2013, 97(9), 3885–3892. doi:10.1007/s00253-012-4681-0. [Crossref] [PubMed]

34. Patricia, G; Cruz, Andrew M. Fribley; Justin R. Miller; Martha J. Larsen; Pamela J. Schultz; Renju T. Jacob; Giselle Tamayo-Castillo; Randal J. Kaufman and Sherman, D.H. Novel Lobophorins inhibit Oral Cancer Cell Growth and Induce Atf4 and chop-Dependent Cell Death in Murine Fibroblasts. ACS Medicinal Chemistry Letters. 2015, 6(8), 877–881. doi:10.1021/acsmedchemlett.5b00127. [Crossref] [PubMed]

35. Zhang, C; Ding, W; Qin, X; and Ju, J. Genome Sequencing of *Streptomyces olivaceus* SCSIO T05 and Activated Production of Lobophorin CR4 via Metabolic Engineering and Genome Mining CR4. Marine Drugs. 2019, 17(10), 1–11. doi:10.3390/md17100593. [Crossref] [PubMed]

36. Luo, M; Tang, L; Dong, Y; Huang, H; Deng, Z; Sun, Y. Antibacterial natural products lobophorin L and M from the marine-derived *Streptomyces* sp. 4506. Nat. Prod. Res. 2021, 35(24), 1–7. doi:10.1080/14786419.2020.1797730. [Crossref] [PubMed]

37. Li, X; Yi-min, CH; Ya-xin, H; Guo-jian, Z; Tian-jiao, Z; Ji-xing, P; Dehai L.C.Q. Isolation and structure elucidation of lobophorin N from the South China Sea-derived *Streptomyces* sp. HDN1844000. J. Antibiot. 2021, 40. doi:1002-3461(2021)05-009-07. [Crossref]

38. Nobutaka Takahashi; Satoshi Miyam oto, Rinpei Mori; Saburo Tamura. Isolation and Physiological Activities of Piericidin A, A Natural Insecticide Produced by *Streptomyces*. Agric. Biol. Chem. 1963, 27, 576–582. doi:10.1080/00021369.1963.10858144. [Crossref]

39. Nobutaka Takahashi, Akinori Suzuki, Satoshi Miyamoto and Saburo Tamura. Structure of piericidin B and stereochemistry of piericidins. Tetrahedron Lett. 1967, 21, 1961–1964. [Crossref]

40. Hiroshi Nishioka; Tsutomu Sawa; Kunio Isshiki, Yoshikazu Takahashi, Naganawa, H; Matsuda, N; Hattori, S; Hamada, M; Takeuchi K.T. Isolation and structure determination of a novel phosphatidylinositol turnover inhibitor, Piericidin B1 N-Oxide. J. Antibiot. 1991, 44, 1283–1285. [Crossref] [PubMed]

41. Yoichi Hayakawa; Shingo Shirasaki; Takashi Kawasaki; Yoshihide Matsuo; Kyoko Adachi Y.S. Structures of New Cytotoxic Antibiotics, Piericidins C7 and C8. J Antibiot. 2007, 60(3), 201–203. [Crossref] [PubMed]

42. Yoshida S; Nagao Y; Watanabe A; Takahashi N. Structure-activity relationship in piericidins inhibitors on the electron transport system in mitochondria. Agric Biol Chem. 1980, 44(12), 2921–2924. doi:10.1080/00021369.1980.10864438. [Crossref]

43. Shaaban, K.A; Helmke, E; Kelter, G; Fiebig, H.H; Laatsch, H. Glucopiericidin C: A cytotoxic piericidin glucoside antibiotic produced by a marine *Streptomyces* isolate. Journal of Antibiotics. 2011, 64(2), 205–209. doi:10.1038/ja.2010.125. [Crossref] [PubMed]

44. Jun-ya Ueda; Takushi Togashi; Susumu Matukura; Aya Nagai; Takuji Nakashima; Hisayuki Komaki; Kozue Anzai; Shigeaki Harayama; Takayuki Doi; Takashi Takahashi; Tohru Natsume; Yasutomo Kisu; Naoki Goshima; Nobuo Nomura; Motoki Takagi. A Novel Nuclear Export Inhibitor JBIR-02, a New Piericidin Discovered from *Streptomyces* sp. ML55. J. Antibiot. 2007, 60(7) 459–462. [Crossref] [PubMed]

45. Wang, M; Carver, J.J; Phelan, V.V; Sanchez, L.M; Garg, N; Peng, Y.,… & Bandeira, N. Sharing and community curation of mass spectrometry data with Global Natural Products Social Molecular Networking. Nature biotechnology. 2016, 34(8), 828–837. doi:10.1038/nbt.3597. [Crossref] [PubMed]

